# *Fas2^EB112^*: A Tale of Two Chromosomes

**DOI:** 10.1101/2024.01.03.574100

**Authors:** Tara M. Finegan, Christian Cammarota, Oscar Mendoza Andrade, Audrey M. Garoutte, Dan T. Bergstralh

## Abstract

The cell-cell adhesion molecule Fasciclin II (Fas2) has long been studied for its evolutionarily-conserved role in axon guidance. It is also expressed in the follicular epithelium, where together with a similar protein, Neuroglian (Nrg), it helps to drive the reintegration of cells born out of the tissue plane. Remarkably, one Fas2 protein null allele, *Fas2^G0336^*, demonstrates a mild reintegration phenotype, whereas work with the classic null allele *Fas2^EB112^* showed more severe epithelial disorganization. These observations raise the question of which allele (if either) causes a *bona fide* loss of Fas2 protein function. The problem is not only relevant to reintegration but fundamentally important to understanding what this protein does and how it works: *Fas2^EB112^* has been used in at least 37 research articles, and *Fas2^G0336^* in at least three. An obvious solution is that one of the two chromosomes carries a modifier that either suppresses (*Fas2^G0336^*) or enhances (*Fas2^EB112^*) phenotypic severity. We find not only the latter to be the case, but identify the enhancing mutation as *Nrg^14^*, also a classic null allele.

## Introduction

The *Drosophila* Immunoglobulin-superfamily cell adhesion molecules Fasciclin II (Fas2, orthologous to vertebrate NCAM) and Neuroglian (Nrg, orthologous to vertebrate L1-CAM), were identified in the developing nervous system, where they localize along fasciculating axons (review (Harden, Wang, and Krieger 2016)). Functional studies have made extensive use of two experimentally-generated null alleles. *Fas2^EB112^*, the first Fas2 allele, was made using imprecise excision of a P-element, resulting in a 1.7kb deletion on the X chromosome that is thought to include the *Fas2* transcriptional start site (Grenningloh, Rehm, and Goodman 1991). *Nrg^14^* (previously called *l(1)RA35*), which was generated using X-ray mutagenesis, is an inversion that disrupts the *Nrg* gene sequence (Lefevre 1981; Hall and Bieber 1997). Early studies demonstrated that both proteins help to drive axon guidance and also that this function is evolutionarily conserved (review (Araújo and Tear 2003)). While their roles in the nervous system have received the most attention, Fas2 and Nrg have also been studied in a variety of developing epithelial tissues, including imaginal discs (Mao and Freeman 2009), Malphigian (renal) tubules (Halberg et al. 2016), the intestine (Resnik-Docampo et al. 2017), the follicular epithelium (Szafranski and Goode 2007; Gomez, Wang, and Riechmann 2012; Fic et al. 2021), and trachea (Neuert et al. 2020) (review (Finegan and Bergstralh 2020)).

One function shared by Fas2 and Nrg is epithelial cell reintegration. Many epithelial cells undergo a change in position during mitosis. Interkinetic nuclear migration (INM), which is the apical-directed movement of the cell nucleus prior to division, is primarily studied in pseudostratified tissues (review (Spear and Erickson 2012)), but apical-directed mitotic cell movement is also evident in simple cuboidal and columnar epithelia, including the mammalian small intestine and ureteric bud (Packard et al. 2013; Sauer 1937; McKinley et al. 2018). In the follicular epithelium, which is cuboidal, roughly half of all cells are born lacking an attachment to the basement membrane (Bergstralh, Lovegrove, and St Johnston 2015). Cells born apical to the plane of the tissue must subsequently incorporate (review (Wilson and Bergstralh 2017)), and this process is mediated by Fas2, Nrg, and to a lesser extent by another IgCAM called Fasciclin III (Fas3) (Bergstralh, Lovegrove, and St Johnston 2015; Cammarota et al. 2020). Genetic disruption of these molecules in the follicular epithelium allows for reintegration to fail, leading to the appearance of apically-positioned cells (Bergstralh, Lovegrove, and St Johnston 2015). We term this phenotype “popping out.”

The functional relationship between Nrg and Fas2 is somewhat perplexing (Williams and Lough 2020). Only about one in 150 cells are popped-out in large mitotic clones (defined as >60% of the follicular epithelium) of either *Fas2^G0336^* (protein null) or *Nrg^14^* (Bergstralh, Lovegrove, and St Johnston 2015; Cammarota et al. 2020). This number increases approximately tenfold in tissue mutant for both alleles (Cammarota et al. 2020). A simple model to explain these findings is that reintegration depends on a total amount of adhesion to which Fas2 and Nrg both contribute. Consistent with this, disruption of either molecule can be largely - though not completely - rescued by additional expression of the other, indicating that they are mostly functionally interchangeable (Cammarota et al. 2020). The same effect is observed for guidance of ocellar pioneer axons in flies (Kristiansen et al. 2005) and is likely to be a conserved feature for the vertebrate orthologs (Kristiansen et al. 2005).

In studying the role that Fas2 plays in reintegration, we encountered an experimental puzzle. Whereas we observed a relatively mild phenotype with *Fas2^G0336^*, another group reported severe tissue disorganization in *Fas2^EB112^* mutant follicular epithelium (Szafranski and Goode 2007). These observations raise the question of why two *Fas2* null alleles have apparently different phenotypic severity, and we address that question here.

## Results

### Two *Fas2^EB112^* chromosomes with different phenotypic severity

*Fas2^EB112^* mutant flies were obtained from the Bloomington Drosophila Stock Center in Indiana, USA (BDSC 36284). These flies were deposited into the Bloomington Drosophila Stock Center in September 2011 and are the only version of *Fas2^EB112^* available from a fly stock depository.

In addition to the *Fas2^EB112^* mutant allele, the X chromosome also includes a site-specific recombination sequence, FRT101, which allows for the generation of mitotic clones. Previous work made use of this technique in the follicular epithelium, a simple monolayer, and found a strong mutant phenotype; mutant tissue is characterized by apically-mispositioned cells that are readily detected (Szafranski and Goode 2007). We repeated these experiments and found the same effect (Supplemental Figure 1A). These results contrast with our own previous work in this tissue. *Fas2^G0336^* mutant clones likewise demonstrate apically-mispositioned cells (henceforth “popped-out,” in agreement with our earlier work), but these events are rare (∼ 1 in 150 cells) (Cammarota et al. 2020). Both *Fas2^EB112^* (Grenningloh, Rehm, and Goodman 1991) and *Fas2^G0336^* are thought to be protein null (Bergstralh, Lovegrove, and St Johnston 2015).

To investigate this difference, we obtained *Fas2^EB112^*mutant flies used in a study performed in the lab of Viet Riechmann in Mannheim, Germany (Gomez, Wang, and Riechmann 2012). The Mannheim *Fas2^EB112^* chromosome makes use of FRT19A, as did our previous cell reintegration studies with *Fas2^G0336^* (Bergstralh, Lovegrove, and St Johnston 2015; Cammarota et al. 2020). Following our established procedure (Bergstralh, Lovegrove, and St Johnston 2015; Cammarota et al. 2020), we measured popping out in follicular epithelia that were >60% homozygous mutant (Figure 1A). Clones made using the Mannheim chromosome had a mild phenotype (∼2 popped-out cells per egg chamber) that is similar to, if slightly weaker than, our previous results using the *Fas2^G0336^* allele (Figure 1A,C) (Bergstralh, Lovegrove, and St Johnston 2015; Cammarota et al. 2020).

**Figure 1:**
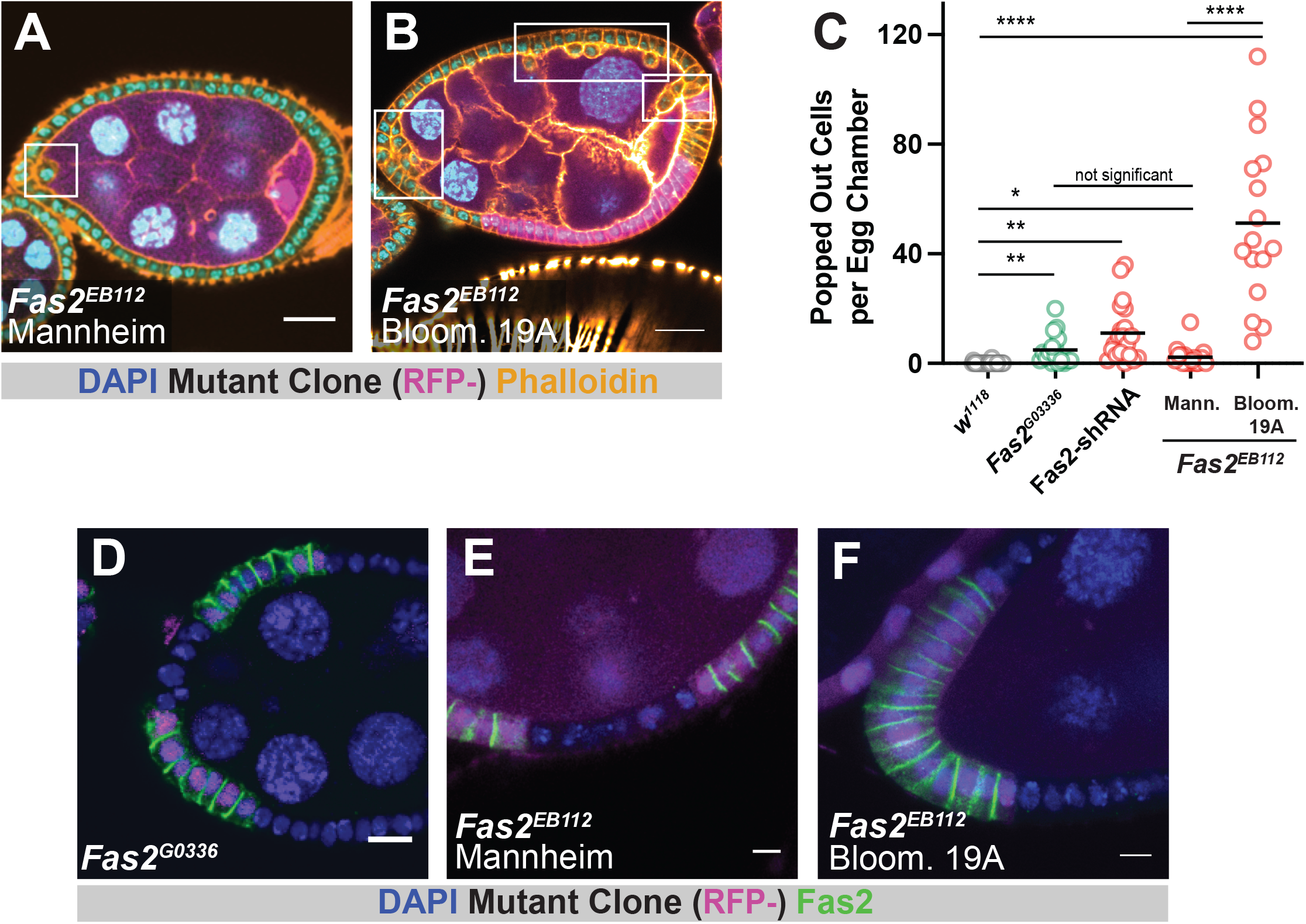
Two *Fas2^EB112^*chromosomes demonstrate different phenotypic severity. **A and B)** Follicle epithelium mutant for Mannheim *Fas2^EB112^* demonstrates rare popping out, whereas Bloom. 19A *Fas2^EB112^* mutant tissue has many popped-out cells. Mutant tissue is marked by the absence of RFP (in magenta). Scale bars = 20μM. **C)** Quantification of popping out shows that the Bloom. 19A *Fas2^EB112^* mutant is significantly more severe than other Fas2-disrupting conditions. **D, E, F)** Fas2 immunoreactivity (measured with the 1D4 antibody) is lost from *Fas2^G0336^*and *Fas2^EB112^* clones. Scale bars = 5μM. Significance was determined using an unpaired t-test with Welch’s correction. In this and subsequent figures, significance is indicated as follows: *p* < 0.05 (*), 0.01 (**), < 0.001 (***), < 0.0001 (****).

We also disrupted Fas2 by expressing UAS-Fas2-shRNA under control of the follicle-cell driver Traffic jam-GAL4. Whereas the mutant analyses are based on large mitotic clones (>60% of the tissue), the knockdown should impact Fas2 protein expression in nearly all of the tissue. Consistent with this, Fas2 knockdown caused the appearance of more popped-out cells (∼11 popped-out cells per egg chamber) than observed using either *Fas2^G0336^*or the Mannheim *Fas2^EB112^* chromosome (Figure 1C), but these egg chambers did not show the severe phenotype associated with the Bloomington chromosome. Together, these results show that the loss of Fas2 is insufficient to explain the severe phenotype observed using the Bloomington *Fas2^EB112^*chromosome.

Because the popping-out phenotype is associated with homozygous lethal mutations, which are investigated using clonal analysis in our system, we recombined the Bloomington chromosome with FRT19A. One advantage to doing this is that it tests whether the difference in flippase recognition target position between the Bloomington and Mannheim *Fas2^EB112^* chromosomes (FRT101 vs. FRT19A, respectively) explains the difference in phenotypic severity.

Our recombination strategy took advantage of the fact that the FRT101 transgenic insertion includes *mini-white,* while the FRT19A insertion does not. We isolated flies without obvious eye coloration conferred by *mini-white* and thereby ensured that recombination took place to the left of the FRT101 insertion, which is at cytological position 14AB (roughly 16MB from the start of the X chromosome).

Through recombination we generated two homozygous lethal FRT19A chromosomes (Supplemental Figure 1B). The first of these (henceforth “*Fas2^EB112^*Bloom. 19A”) resembles the Bloomington *Fas2^EB112^* chromosome: 1) Like *Fas2^G0336^* and Mannheim *Fas2^EB112^*, mutant clones generated using the *Fas2^EB112^* Bloom. 19A chromosome lack Fas2 expression as measured with the 1D4 antibody (Figure 1D-F). 2) Significantly more popped-out cells are observed in tissue homozygous for the *Fas2^EB112^* Bloom. 19A chromosome, indicating that FRT101 does not explain the difference in severity (Figure 1B,C). Taken together, our results indicate that the original Bloomington *Fas2^EB112^*and the derived *Fas2^EB112^* Bloom. FRT19A chromosomes carry a secondary mutation that enhances popping out.

### *E(Fas2) ^mut^* enhances popping out but does not affect spindle orientation

The second chromosome that we generated through recombination did not demonstrate decreased anti-Fas2 immunoreactivity or the presence of popped-out cells (Figure 2A,B), indicating that the original Bloomington chromosome has at least one other lethal mutation besides *Fas2^EB112^*. We considered the possibility that this chromosome might harbor the phenotype-enhancing mutation. To test this, we generated mitotic clones in tissue expressing UAS-Fas2-shRNA. We found that the number of popped-out cells increased significantly over Fas2-knockdown alone (Figure 2B, Supplemental Movie 1). We therefore provisionally identify the mutation on our second chromosome as *Enhancer of Fas2^mut^* (*E(Fas2)^mut^*).

**Figure 2:**
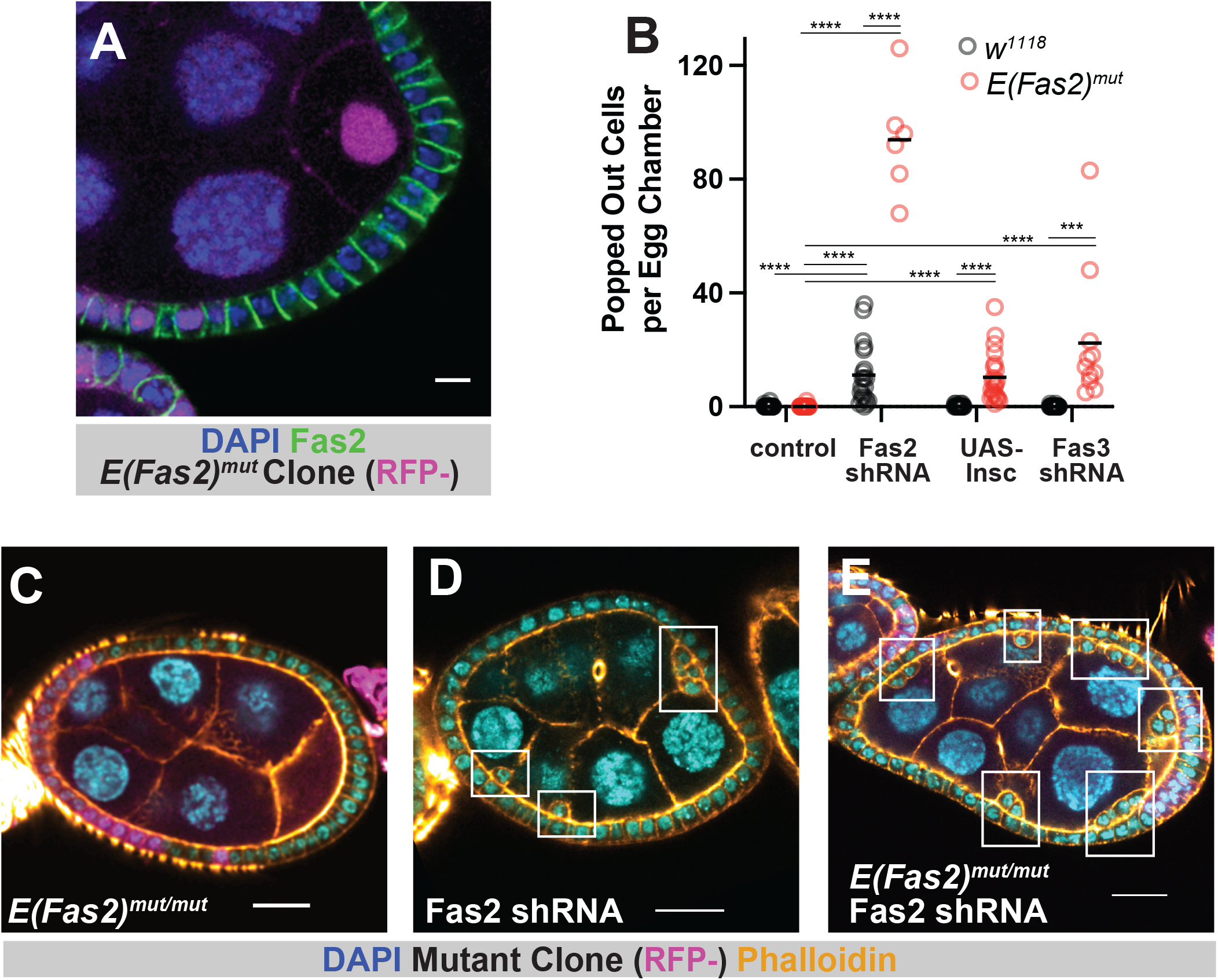
Characterization of *E(Fas2)^mut^*. **A)** Fas2 is detected at follicle cell-cell borders in *E(Fas2)^mut^* tissue. Scale bar = 5μM. **C, D, E)** Representative images showing the enhancement of popping out in Fas2-shRNA tissue. Scale bars = 20μM.

Our earlier work defined two pathways towards enhancement of the popping-out phenotype caused by *Fas2* disruption. The first of these is misorientation of the mitotic spindle, which leads to misoriented cell divisions, and the second is disruption of another lateral adhesion protein. Epithelial cells typically orient their spindles perpendicular to the apical-basal axis so that new cells are born roughly along the plane of the tissue (reviewed in (Bergstralh, Dawney, and St Johnston 2017)). The popping out phenotype is enhanced by ectopic expression of Inscuteable, a protein that is normally expressed in neural progenitor cells (neuroblasts) in the developing nervous system (Bergstralh, Lovegrove, and St Johnston 2015; Kraut et al. 1996). When it is expressed in the follicular epithelium, Inscuteable can reorient spindles so that they are parallel to the apical-basal axis, causing some daughter cells to be born outside the tissue plane (Bergstralh, Lovegrove, and St Johnston 2015; Neville et al. 2023). This manipulation does not cause tissue disorganization by itself because the misplaced cells reintegrate. However, it does increase the number of popped-out cells if the reintegration mechanism is impaired, as it is in *Fas2* or *Nrg* mutant tissue.

We asked whether *E(Fas2)^mut^*, like Inscuteable, impacts spindle orientation. It does not (Supplemental Figure 2A, B), and this finding rules out the first known pathway to phenotypic enhancement. We therefore considered whether *E(Fas2)^mut^* uses the second pathway, namely disruption of another lateral adhesion protein besides Fas2. As a first test of this possibility we asked whether ectopic Inscuteable expression increases the number of popped-out cells in *E(Fas2)^mut^* tissue. We find that it does (Figure 2B). Together, these results indicate that *E(Fas2)^mut^* behaves like mutations that impair lateral adhesion molecules.

Three IgCAMs – namely, Fas2, Nrg, and Fas3 – are known to participate in reintegration. Fas3 plays a smaller role than either Nrg or Fas2; loss of Fas3 function using a strong knockdown exacerbates popping out in Nrg or Fas2 mutants but does not itself lead to popping out (Cammarota et al. 2020). The same is true for *E(Fas2)^mut^*and we therefore asked whether *E(Fas2)^mut^* impacts Fas3 localization or expression in the follicular epithelium. We do not see loss of Fas3 immunoreactivity at cell-cell borders in *E(Fas2)^mut^* tissue (Supplemental Figure 2C), suggesting that *E(Fas2)^mut^* does not disrupt Fas3 function. We also tested for genetic interaction and found that the combination of *E(Fas2)^mut^* and *Fas3* knockdown leads to the appearance of popped-out cells (Figure 2B). These results show that *E(Fas2)^mut^* does not work through disruption of Fas3.

### *E(Fas2)^mut^* causes disruption of Neuroglian

There are nine Nrg mRNAs currently annotated (FlyBase). The protein coding sequences for all nine mRNAs are identical over a span of 3,372 nucleotides (1,124 AAs) encoded by the same six exons, after which point sequences diverge into three groups based on the terminal (seventh) protein-coding exon (Supplemental Figure 3A). The terminal exon used in the first group (mRNAs A, C, D, F, and H) encodes 17 amino acids, and these include the highly-conserved FIGQY subsequence that is A) a hallmark of Nrg proteins across species (reviewed in (Michael Hortsch, Nagaraj, and Godenschwege 2009)) and B) implicated in epithelial cell reintegration (Cammarota et al. 2020). These mRNAs encode the Nrg^167^ protein isoform that is expressed outside of the nervous system (M Hortsch et al. 1990). The second group (Isoforms B, E, and I) uses a terminal exon that encodes 80 amino acids, also including a FIGQY subsequence. These mRNA isoforms encode the Nrg^180^ protein expressed in the nervous system (M Hortsch et al. 1990). The final group includes only one isoform (G) and is unique in that it does not have the FIGQY subsequence.

We generated a rabbit polyclonal antibody using the sequence K1160-V1302 (Nrg^180^) as the antigen. This sequence includes 64 AAs shared between all isoforms and an additional 10 that are highly similar between Nrg^167^ and Nrg^180^. The antibody detects signal at cell-cell borders in proliferative-stage follicle epithelia, consistent with prior work (Wei, Hortsch, and Goode 2004). Immunoreactivity is lost in *Nrg^14^* (null) mutant clones, demonstrating specificity (Supplemental Figure 3B).

Neither our antibody nor a previously generated mouse monoclonal antibody (Bieber et al. 1989) detect immunoreactivity at cell-cell borders in *e(Fas2)^mut^*homozygous clones (Figure 3A-C). Consistent with this result, immunoreactivity is not observed in *Fas2^EB112^* Bloom. 19A clones (Figure 2E). It is, however, readily apparent in *Fas2^EB112^* Mannheim clones. We conclude that *e(Fas2)^mut^* causes the disappearance of Nrg protein from cell-cell borders.

**Figure 3:**
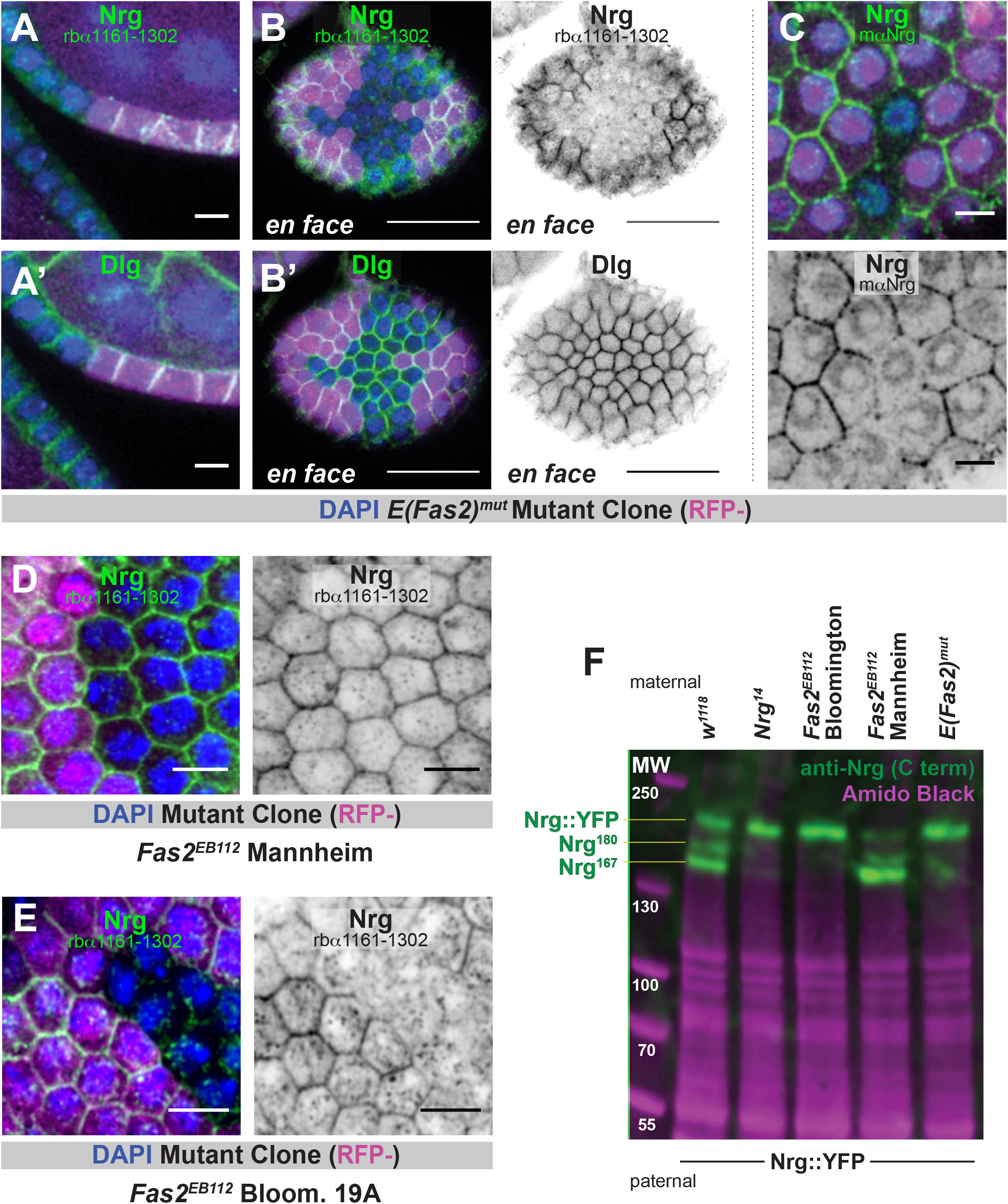
Nrg is not expressed in *E(Fas2)^mut^* tissue. **A and B)** Anti-Nrg immunoreactivity is lost from follicle cell-cell borders in *E(Fas2)^mut^* clones (marked by the absence of RFP). These images were generated using a rabbit polyclonal antibody that recognizes a C-terminal stretch of Nrg. Two views, sagittal (A) and *en face* (B) are shown. Discs large (Dlg), which localizes to cell-cell borders in a similar manner to Nrg, is used as a control. Scale bars = 5μM (A) or 20μM (B). **C)** A mouse monoclonal anti-Nrg antibody also fails to detect signal at follicle cell-cell borders in *E(Fas2)^mut^* clones. Scale bars = 5μM. **D and E)** Anti-Nrg immunoreactivity is retained at cell-cell borders in Mannheim *Fas2^EB112^* tissue (D) but lost in Bloom. 19A *Fas2^EB112^* mutant tissue (E). Scale bars = 5μM. **F)** Immunoblotting reveals that *E(Fas2)^mut^* and Bloomington *Fas2^EB112^*chromosomes do not contribute expression of Nrg^167^ and Nrg^180^ protein isoforms, whereas the Mannheim *Fas2^EB112^* chromosome does. *Nrg^14^* is used as a negative control. Expression of Nrg::YFP is stronger when Nrg^167^ and Nrg^180^ are lost. Amido black staining reveals total protein and is therefore a loading control. Significance was determined using an unpaired t-test.

A straightforward possibility is that *e(Fas2)^mut^* prevents Nrg protein expression. We performed a western blot to test this, using lysate from whole adult female flies. To generate these flies we crossed females from multiple fly lines of interest to males of the genotype *Nrg::YFP* (Lowe et al. 2014). The logic behind this experimental design is that Nrg::YFP (inherited from the father) can be easily distinguished from untagged Nrg (inherited from the mother) on the basis of protein size; the yellow fluorescent protein (YFP) tag adds approximately 28 kDa. We note that occasional nondisjunction of the sex chromosomes was noticed in our experiments with the *Fas2^EB112^* Bloomington chromosome. This issue did not affect our protein expression results or subsequent experiments because genotypes could be easily distinguished on the basis of eye shape and color (Supplemental Figure 3C).

Three bands are observed in the positive control lane (*w^1118^*) (Figure 3F). The lower two correspond in position to Nrg^167^ and Nrg^180^. We identify the highest band as Nrg::YFP; it is also observed in the negative control lane (*Nrg^14^*) (Figure 3F) and when the same lysates are probed using anti-GFP antibody (not shown). The finding that only one band is observed at this highest molecular weight indicates expression of either Nrg^180^::YFP or Nrg^167^::YFP but not both. Based on size, we expect the former. The Nrg::YFP line (aka *Nrg^CPTI001714^*) was generated as part of the Cambridge Protein Trap Insertion screen, in which an artificial exon encoding YFP was randomly inserted into intron sequences. In this case, the artificial exon is inserted at X chromosome position 8,549,429 bp, inside the final *Nrg* intron and < 1kb from the start of the *Nrg^167^* terminal exon (Supplemental Figure 3A). We speculate that the insertion interferes with splicing, causing the *Nrg^167^* exon to be skipped.

Nrg^167^ and Nrg^180^ are not observed in the *Fas2^EB112^* Bloomington or *E(Fas2)^mut^* lanes (Figure 3F). Both proteins are, however, apparent in the *Fas2^EB112^* Mannheim lane. We also observed that the intensity of the Nrg::YFP (highest) band increased in the negative control, the *Fas2^EB112^* Bloomington, and the *E(Fas2)^mut^* lanes (Figure 3F and Supplemental Figure 3D). Together, these results indicate that *E(Fas2)^mut^* prevents the expression of Nrg^167^ and Nrg^180^. They also show that the loss of Nrg expression from one chromosome is compensated by enhanced expression from the other.

### *E(Fas2)^mut^* is *Nrg^14^*

Loss of Nrg function is associated with lethality (Hall and Bieber 1997) and with the enhancement of reintegration failure in *Fas2* mutants (Cammarota et al. 2020). Both phenotypes are also observed for *E(Fas2)^mut^*, and we therefore asked whether they could be rescued by ectopic Nrg expression. To test for this we took advantage of two genomic rescue strategies: 1) a large transgenic insertion that includes *Nrg* (with its promoter) on the second chromosome (Enneking et al. 2013) and 2) a Y chromosome to which a duplication of the genomic region encoding *Nrg* has been attached (Cook et al. 2010). Expression of *Nrg* using either strategy allows for the viability of *E(Fas2)^mut^* / Y and *Nrg^14^* / Y males but not *Fas2^EB112^* Mannheim / Y males (Figure 4A and Supplemental Figure 4A). Furthermore, expression of *Nrg* from the second chromosome rescues the popping out phenotype observed in *Fas2^EB112^* Bloom. FRT19A mutant tissue (Figure X). Consistent with this, we find that *E(Fas2)^mut^* does not increase the number of popped-out cells in *Nrg* knockdown tissue (Figure 4B). Taken together, these results show that Nrg disruption can explain both lethality and the enhancement of reintegration failure caused by *E(Fas2)^mut^*.

**Figure 4:**
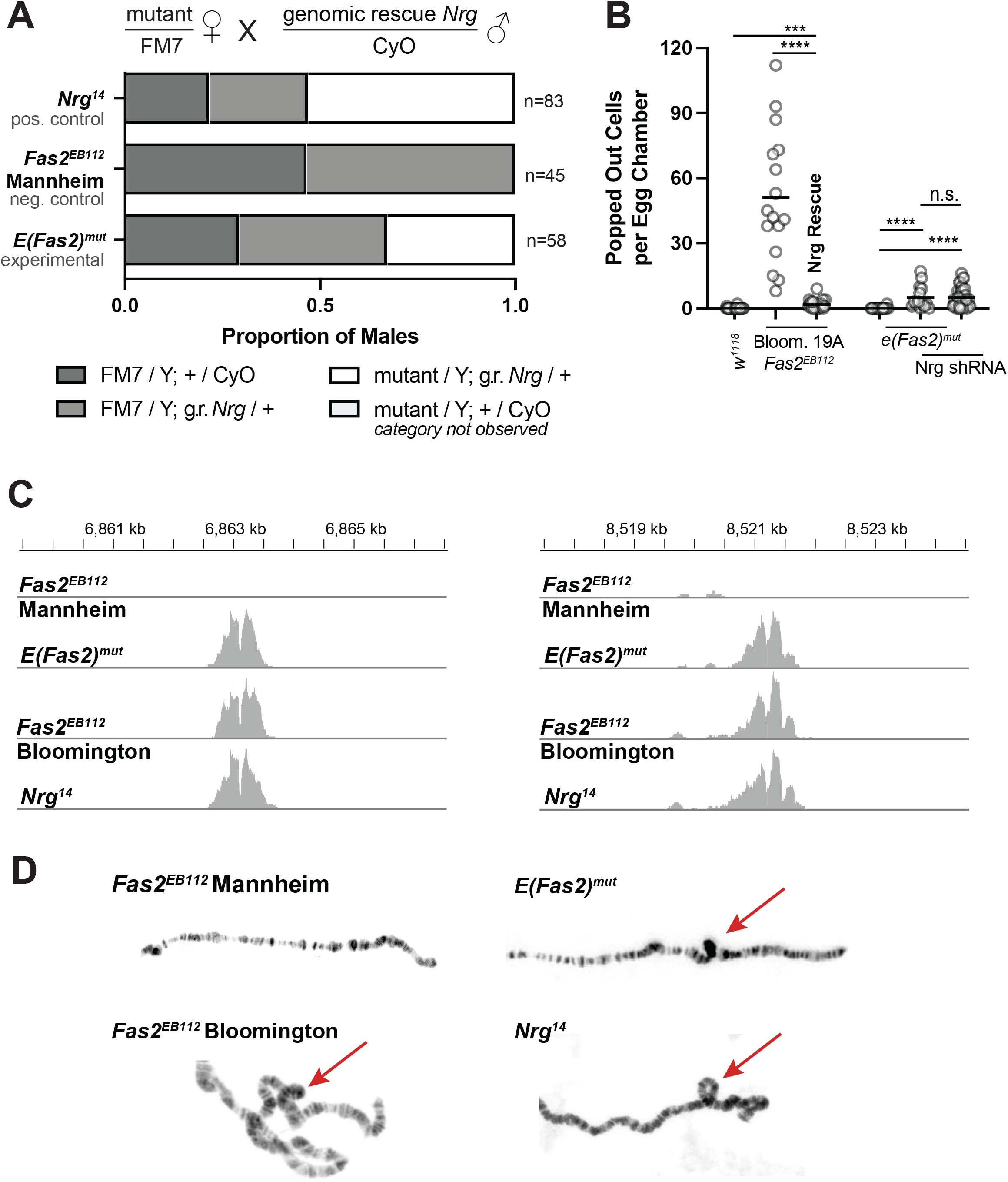
*E(Fas2)^mut^*is *Nrg^14^*. **A)** Expression of Nrg from the second chromosome rescues the viability of *E(Fas2)^mut^* male flies. Balanced mutant females were crossed to males carrying an Nrg genomic rescue insertion on the second chromosome. This strategy allowed for the appearance of *Nrg^14^* and *E(Fas2)^mut^* male progeny. **B)** Expression of Nrg from the second chromosome rescues popping out in Bloom. 19A *Fas2^EB112^* mutant tissue. (The Bloom. *Fas2^EB112^*data in this figure is also shown in Figure 1C.) Additionally, Nrg knockdown causes the appearance of popped-out cells but this phenotype is not enhanced by *E(Fas2)^mut^*. **C)** Cumulative plot of broken sequencing reads and their position along the chromosome. Sequencing was performed at 100X coverage and the Y-axis scale is set at 0-100. The X axis (position along the X chromosome) is shown. The central divots indicate small deletions described in the text. **D)** Polytene X chromosomes from female larvae with one X chromosome from the indicated genotype and the other from a *w^1118^*male. Loops that indicate the common inversion are highlighted with a red arrow.

Our findings raise the obvious question of whether *E(Fas2)* is *Nrg.* To answer this we first performed Illumina short-read sequencing on flies with the following chromosomes: *E(Fas2)^mut^*, Mannheim *Fas2^EB112^*, Bloomington *Fas2^EB112^*, and *Nrg^14^*. The deletion that disrupts *Fas2* in the two EB112 lines is readily identified at 4,205,492 - 4,207,081 bp, and this finding is both consistent with and extends earlier work (Grenningloh, Rehm, and Goodman 1991). Based on single nucleotide polymorphisms, we find that the Bloomington *Fas2^EB112^* and *E(Fas2)^mut^* chromosomes are highly similar between nucleotides ∼4,790 kb and ∼13,180 kb (using dm6 as the reference genome), indicating that these are the approximate positions at which two crossovers were resolved when the latter chromosome was generated (Supplemental Figure 4B). The *eFas2* locus is therefore within that span. So is *Nrg*.

Sequencing reads for the *E(Fas2)^mut^*, Bloomington *Fas2^EB112^*, and *Nrg^14^* chromosomes were broken at or near a small deletion interrupting the coding sequence of *CG14434* (nts 6,863,120-125) and continued at or near another deletion within the first intron of *Nrg* (nts 8,521,186-191) (Figure 4C). These reads indicate an inversion between cytological positions 6E and 7F1, which is the *Nrg^14^* allele. We confirmed this finding by examining polytene chromosomes, in which inversions are revealed as loops. When paired with a control chromosome (*w^1118^*), the *E(Fas2)^mut^*, Bloomington *Fas2^EB112^*, and *Nrg^14^* chromosomes all exhibited loops at the location indicated by our sequencing data, whereas the Mannheim *Fas2^EB112^*chromosome does not (Figure 4D). Together, these results show that *E(Fas2)^mut^* is *Nrg^14^*.

## Discussion

Although similar, our *E(Fas2)^mut^* and *Nrg^14^* mutants are not phenotypically identical. *E(Fas2)^mut^*has a weaker phenotype: we do not observe popped-out cells in this tissue and we find fewer popped-out cells when this allele is combined with Inscuteable or Fas3-shRNA than in our previous work with *Nrg^14^*. Similarly, Mannheim *Fas2^EB112^* has a lower average number of popped-out cells than *Fas2^G0336^* (Figure 1C), though this difference is not significant. These results suggest that other factors modulate the expression or activity of reintegration factors. Because popped-out cells have been observed in other *Nrg*-disruption conditions, namely knockdown (Bergstralh, Lovegrove, and St Johnston 2015) and the temperature sensitive allele I(1)B4 (Wei, Hortsch, and Goode 2004), we suspect that *E(Fas2)^mut^* is suppressed. We have used our *E(Fas2)^mut^* and *Nrg^14^* chromosomes in several genetic backgrounds (with respect to autosomes) and found the same results, suggesting that modulation is due to differences on the X chromosome. The short read sequencing we performed permits comparison of single nucleotide polymorphisms and small indels on this chromosome, but we do not identify any of these changes as obvious candidates for modulating reintegration.

Additional results also suggest that analysis of reintegration factors may be more complicated than anticipated by our own previous work. We find that loss of Nrg protein expression from one gene copy is compensated by increased expression from the other (Figure 3F and Supplemental Figure 3C), meaning that heterozygosity should not be expected to substantially impact function. Whether this mechanism or a similar one also promotes expression of other reintegration factors is unknown, though we did not see obviously increased expression of either Fas2 or Fas3 in *E(Fas2)^mut^*clonal tissue.

Earlier work shows that reintegration relies on a combination of adhesion factors that act in partial redundance. The observation that Nrg expression is strictly regulated indicates adds yet another layer of robustness, and thereby underlines the importance of this process.

## Acknowledgments

We are grateful to Juan Manuel Gomez; Jeff Sekelsky; the labs of Mark Peifer, Scott Williams, and Holly Lovegrove; Jim Birchler and other members of the University of Missouri Fly Club; and members of the Finegan-Bergstralh lab for their questions and comments. This work was supported by an NSF CAREER award (PI: Bergstralh) and NIH Grant R01GM125839 (PI: Bergstralh).

## Competing Interests

The authors declare no competing interests.

---

**Supplemental Figure 1: A)** A severe popping out phenotype is observed in homozygous mutant clones generated using the Mannheim *Fas2^EB112^*chromosome. Scale bars = 20μM. **B)** A schematic showing how the two lethal chromosomes were generated through recombination. The original Bloomington *Fas2^EB112^* chromosome is represented in red and the FRT19A chromosome with which it was recombined is in blue. Purple represents sequence that may have come from either of these. At this point in the study we could not distinguish whether the *E(Fas2)^mut^* chromosome was the product of a single crossover event to the left of the *Fas2* locus or two crossovers to the right (see Supplemental Figure 4).

**Supplemental Figure 2: A and B)** Mitotic spindle angles are parallel to the tissue plane in *E(Fas2)^mut^* tissue. Representative image in (A) and quantification in (B). **C)** Fas3 immunoreactivity at follicle cell-cell borders is retained in *E(Fas2)^mut^* tissue. Scale bars in (A) and (C) = 5μM.

**Supplemental Figure 3: A)** Diagram illustrating splicing variants that generate different Nrg protein isoforms. The position of the YFP insertion (*Nrg^CPTI001714^*) is also shown. **B)** Immunostaining confirms the specificity of the anti-Nrg antibody. Scale bar = 20μM. **C)** Occasional nondisjunction of the sex chromosomes results in XXY and XO flies, which can be easily distinguished (and therefore excluded) by eye shape and eye color. **D)** Quantification of Nrg::YFP protein expression (as measured by immunoblot band intensity) across samples shows that it is enhanced in samples lacking Nrg^167^ and Nrg^180^ (related to Figure 2F). Intensity was measured in four immunoblots based on two lysate preparations. Error bars represent standard deviation.

**Supplemental Figure 4: A)** Expression of Nrg from the Y chromosome rescues the viability of *E(Fas2)^mut^* male flies. Balanced mutant females were crossed to males that have Y chromosomes with a duplication of the X chromosome that includes *Nrg*. This strategy allowed for the appearance of *Nrg^14^* and *E(Fas2)^mut^*male progeny. **B)** Single nucleotide polymorphisms (compared to the dm6 reference genome) reveal sequence similarity between the Mannheim *Fas2^EB112^* chromosome and the Bloomington *Fas2^EB112^* chromosome, and also similarity between the Bloomington *Fas2^EB112^* chromosome and the *E(Fas2)^mut^* chromosome. The latter comparison reveals extensive similarity over a region that includes *Nrg*. Both nucleotide position and a cytological map are shown for reference.

**Supplemental Movie 1:** Genetic disruption of both *Fas2* (knockdown driven by Traffic jam-GAL4) and *E(Fas2)* (*E(Fas2)^mut^*clones marked by the absence of RFP, in magenta) causes a severe popping out phenotype. The video is a z-axis fly-through of a stack of images spaced 0.5 microns apart. Actin (phalloidin) is revealed in orange and DNA (DAPI) in cyan.

## Materials and Methods

### Reagents

A list of reagents used in this study is found in Table 1.

### *Drosophila* genetics

A list of alleles and transgenes used in this study is found in Table 2. We thank the Transgenic RNAi Project at Harvard Medical School (NIH/NIGMS R01-GM084947) for providing shRNA lines. Ectopic protein expression was accomplished using the UAS-GAL4 system (Brand and Perrimon 1993). Expression was driven by Traffic Jam-GAL4 (Olivieri et al. 2010).

### Mitotic clones

The recombinase (flippase) is under control of a heat shock promoter. Mitotic clones were generated by incubating larvae or pupae at 37°C for two out of every twelve hours over a period of at least two days. Ovaries were dissected from adult flies at least two days after the last heat shock. Flies in which the Gal4-UAS system was used were kept at 29° for at least 48 hours before dissection.

### Misplaced Cell Counting

Quantification of extra-layer cells was performed on Stage 6-8 egg chambers using at least 3 dissections of at least 5 flies each. For analyses of clonal mutants, the number of extra-layer cells was quantified in egg chambers that were at least 60% mutant. Popped-out cells were quantified manually. Each data point reflects the total number of misplaced cells (examined through the entire depth) in an egg chamber. Images are representative sagittal planes.

### Immunostaining

Ovaries were fixed for 15 minutes in 10% Formaldehyde and 0.2% Tween in Phosphate Buffered Saline (PBS-T) and subsequently incubated in blocking solution (10% Bovine Serum Albumin in PBS) for approximately one hour at room temperature. Primary and secondary immunostainings lasted 12 or more hours at 4°C in PBS-T. Three washes of about 5 minutes each in PBS-T were carried out after the primary and secondary stainings. Both primary and secondary antibodies were used at a concentration of 1:150.

### Imaging

Microscopy was performed using a Leica SP5 point scanning confocal (63x/1.4 HCX PL Apo CS oil lens) or Leica SP8 point scanning confocal (63x/1.4 HCX PL Apo CS oil lens). Images were collected with LAS AF. Minor processing (Gaussian blur) was performed using FIJI.

### Sequencing

2X 150-bp paired-end genome sequencing at ∼100X coverage was performed by the University of Missouri Genomics Technology Core using the Illumina NovaSeq platform. Analysis was performed with help from the University of Missouri bioinformatics core. Raw reads were filtered by fastp (Chen et al. 2018). The clean reads were used for mapping and variant calling, which was performed using the Parabricks pbrun germline command with the *Drosophila* reference genome GCF_000001215.4 (NCBI). Variants were subsequently hard-filtered by GATK VariantFiltration following its best practice (Franke and Crowgey 2020). The VCF file *nexus.all-chrs.vars.AN268.R6.share.verboseINFO.vcf.gz* (FlyBase Associated Files, updated 11-27-23), which contains updated *Drosophila melanogaster* genetic variant data, was used to identify reported/common variants. BAM files were visualized using IGV (Robinson et al. 2023).

### Western Blots and Quantification

Whole flies were lysed in the following buffer: 1% Triton X-100, 150 mm NaCl, 20mM Tris-HCl, 1mM EGTA, 1 mM EDTA, plus a protease inhibitor mixture (Roche Applied Science). Samples were resolved by SDS-PAGE and transferred to PVDF. Immunoblots were probed with the anti-Nrg primary antibody for >24 hours at 4°C in PBS-T, washed three times in PBS-T, and probed with secondary for >24 hours at 4°C in PBS-T. Immunoreactivity was visualized using enhanced chemiluminescence. As a control for protein loading, the blot was stained with Amido Black. We quantified band intensity using FIJI as follows: A freehand line drawn through all five bands was used to generate a pixel intensity plot. A corrected intensity for each band was generated by subtracting background signal from that band’s maximum intensity. The data were normalized by dividing each band’s corrected intensity by the mean average corrected intensity of all five bands.

### Statistics Software

Statistical analysis was performed using GraphPad Prism.

### Generation of anti-Neuroglian antibody

The Nrg antibody was designed and generated by ABclonal. Rabbits were injected with a synthetic peptide from *Nrg^180^* sequence 1161-1302. Antisera were collected and affinity purified.

### Polytene chromosomes

Female 3rd instar wandering larvae from a cross of female flies possessing the chromosome of interest over FM7-RFP to *w-* males were selected for an absence of RFP using a fluorescence widefield microscope. Salivary glands were dissected from these larvae in PBS and the fat body removed. Glands were transferred to 100 µl of fresh fixation solution for 1 minute (2% Paraformaldehyde, 45% Acetic Acid in MilliQ water). Glands were then transferred to 7 µl drop of a fresh dilution of 45% Acetic Acid in MillQ water on a coverslip pre-treated with Sigmacote. A poly-L-lysine coated slide was lowered onto the coverslip on top of the glands. Glands were squashed by applying pressure with a gloved finger in a clockwise rotation and then a rubber stopper was used to apply medium force 25 times to the slide wrapped in thick filter paper. Slides were then dipped into liquid nitrogen. Once returned to room temperature, the cover slip was removed using a razor blade and discarded. Once all liquid condensation was evaporated from the slide, 200 µl of Vectashield with DAPI was applied on top of the glands and a new coverslip applied and sealed with nail polish. After at least 3 hrs, DAPI stained glands were imaged using a Leica SP8 confocal microscope using an HC PLAN APO CS2 63/1.40 objective.

### Rescue counts

Crosses were set up as indicated in Figure 4. and maintained at 26°C. The cross (parentals) were transferred into a new vial every 3-4 days. Progeny from at least three of these vials were collected and male genotypes were scored on the basis of phenotypic markers. To account for the possibility that males of different genotypes might eclose at different rates, males were counted until no more flies eclosed from the vial. In the case of the Mannheim *Fas2^EB112^*/ *FM7 X PAC Nrg / CyO* cross, six vials were collected.

## Notes

### Competing Interest Statement

The authors have declared no competing interest.

## References

Araújo, Sofia J, and Guy Tear. 2003. “Axon Guidance Mechanisms and Molecules: Lessons from Invertebrates.” Nature Reviews. Neuroscience 4 (11): 910–22.

Bergstralh, Dan T, Nicole S Dawney, and Daniel St Johnston. 2017. “Spindle Orientation: A Question of Complex Positioning.” Development 144 (7): 1137–45.

Bergstralh, Dan T, Holly E Lovegrove, and Daniel St Johnston. 2015. “Lateral Adhesion Drives Reintegration of Misplaced Cells into Epithelial Monolayers.” Nature Cell Biology 17 (11): 1497–1503.

Bieber, Allan J, Peter M Snow, Michael Hortsch, Nipam H Patel, J Roger Jacobs, Zaida R Traquina, Jim Schilling, and Corey S Goodman. 1989. “Drosophila Neuroglian: A Member of the Immunoglobulin Superfamily with Extensive Homology to the Vertebrate Neural Adhesion Molecule L1.” Cell 59 (3): 447–60.

Brand, A H, and N Perrimon. 1993. “Targeted Gene Expression as a Means of Altering Cell Fates and Generating Dominant Phenotypes.” Development 118 (2): 401–15.

Cammarota, Christian, Tara M. Finegan, Tyler J. Wilson, Sifan Yang, and Dan T. Bergstralh. 2020. “An Axon-Pathfinding Mechanism Preserves Epithelial Tissue Integrity.” Current Biology 30 (24): 5049–5057.e3. 10.1016/j.cub.2020.09.061.

Chen, Shifu, Yanqing Zhou, Yaru Chen, and Jia Gu. 2018. “Fastp: An Ultra-Fast All-in-One FASTQ Preprocessor.” Bioinformatics 34 (17): i884–90. 10.1093/bioinformatics/bty560.

Cook, R. Kimberley, Megan E. Deal, Jennifer A. Deal, Russell D. Garton, C. Adam Brown, Megan E. Ward, Rachel S. Andrade, Eric P. Spana, Thomas C. Kaufman, and Kevin R. Cook. 2010. “A New Resource for Characterizing X-Linked Genes in Drosophila Melanogaster: Systematic Coverage and Subdivision of the X Chromosome With Nested, Y-Linked Duplications.” Genetics 186 (4): 1095–1109. 10.1534/genetics.110.123265.

Enneking, Eva-Maria, Sirisha R Kudumala, Eliza Moreno, Raiko Stephan, Jana Boerner, Tanja A Godenschwege, and Jan Pielage. 2013. “Transsynaptic Coordination of Synaptic Growth, Function, and Stability by the L1-Type CAM Neuroglian.” 11 (4): e1001537.

Fic, Weronika, Rebecca Bastock, Francesco Raimondi, Erinn Los, Yoshiko Inoue, Jennifer L. Gallop, Robert B. Russell, and Daniel St Johnston. 2021. “RhoGAP19D Inhibits Cdc42 Laterally to Control Epithelial Cell Shape and Prevent Invasion.” Journal of Cell Biology 220 (4): e202009116. 10.1083/jcb.202009116.

Finegan, Tara M., and Dan T. Bergstralh. 2020. “Neuronal Immunoglobulin Superfamily Cell Adhesion Molecules in Epithelial Morphogenesis: Insights from Drosophila.” Philosophical Transactions of the Royal Society B: Biological Sciences 375 (1809): 20190553. 10.1098/rstb.2019.0553.

Franke, Karl R., and Erin L. Crowgey. 2020. “Accelerating next Generation Sequencing Data Analysis: An Evaluation of Optimized Best Practices for Genome Analysis Toolkit Algorithms.” Genomics & Informatics 18 (1). 10.5808/GI.2020.18.1.e10.

Gomez, Juan Manuel, Ying Wang, and Veit Riechmann. 2012. “Tao Controls Epithelial Morphogenesis by Promoting Fasciclin 2 Endocytosis.” The Journal of Cell Biology 199 (7): 1131–43.

Grenningloh, G, E J Rehm, and C S Goodman. 1991. “Genetic Analysis of Growth Cone Guidance in Drosophila: Fasciclin II Functions as a Neuronal Recognition Molecule.” Cell 67 (1): 45–57.

Halberg, Kenneth A, Stephanie M Rainey, Iben R Veland, Helen Neuert, Anthony J Dornan, Christian Klämbt, Shireen-Anne Davies, and Julian A T Dow. 2016. “The Cell Adhesion Molecule Fasciclin2 Regulates Brush Border Length and Organization in Drosophila Renal Tubules.” Nature Communications 7 (1): 1–10.

Hall, Stephen G., and Allan J. Bieber. 1997. “Mutations in the Drosophila Neuroglian Cell Adhesion Molecule Affect Motor Neuron Pathfinding and Peripheral Nervous System Patterning.” Journal of Neurobiology 32 (3): 325–40. 10.1002/(SICI)1097-4695(199703)32:3<325::AID-NEU6>3.0.CO;2-9.

Harden, Nicholas, Simon Ji Hau Wang, and Charles Krieger. 2016. “Making the Connection - Shared Molecular Machinery and Evolutionary Links Underlie the Formation and Plasticity of Occluding Junctions and Synapses.” Journal of Cell Science 129 (16): 3067–76.

Hortsch, M, A J Bieber, N H Patel, and C S Goodman. 1990. “Differential Splicing Generates a Nervous System-Specific Form of Drosophila Neuroglian.” Neuron 4 (5): 697–709.

Hortsch, Michael, Kakanahalli Nagaraj, and Tanja A. Godenschwege. 2009. “The Interaction between L1-Type Proteins and Ankyrins - a Master Switch for L1-Type CAM Function.” Cellular & Molecular Biology Letters 14 (1): 57–69. 10.2478/s11658-008-0035-4.

Kraut, R, W Chia, L Y Jan, Y N Jan, and J A Knoblich. 1996. “Role of Inscuteable in Orienting Asymmetric Cell Divisions in Drosophila.” Nature 383 (6595): 50–55.

Kristiansen, Lars V, Emma Velasquez, Susana Romani, Sigrid Baars, Vladimir Berezin, Elisabeth Bock, Michael Hortsch, and Luis Garcia-Alonso. 2005. “Genetic Analysis of an Overlapping Functional Requirement for L1- and NCAM-Type Proteins during Sensory Axon Guidance in Drosophila.” Molecular and Cellular Neurosciences 28 (1): 141–52.

Lefevre, George. 1981. “The Distribution of Randomly Recovered X-Ray-Induced Sex-Linked Genetic Effects in DROSOPHILA MELANOGASTER.” Genetics 99 (3–4): 461. 10.1093/genetics/99.3-4.461.

Lowe, Nick, Johanna S Rees, John Roote, Ed Ryder, Irina M Armean, Glynnis Johnson, Emma Drummond, et al. 2014. “Analysis of the Expression Patterns, Subcellular Localisations and Interaction Partners of Drosophila Proteins Using a pigP Protein Trap Library.” Development 141 (20): 3994–4005.

Mao, Yanlan, and Matthew Freeman. 2009. “Fasciclin 2, the Drosophila Orthologue of Neural Cell-Adhesion Molecule, Inhibits EGF Receptor Signalling.” Development 136 (3): 473–81. 10.1242/dev.026054.

McKinley, Kara L, Nico Stuurman, Loic A Royer, Christoph Schartner, David Castillo-Azofeifa, Markus Delling, Ophir D Klein, and Ronald D Vale. 2018. “Cellular Aspect Ratio and Cell Division Mechanics Underlie the Patterning of Cell Progeny in Diverse Mammalian Epithelia.” eLife 7 (June): 19.

Neuert, Helen, Petra Deing, Karin Krukkert, Elke Naffin, Georg Steffes, Benjamin Risse, Marion Silies, and Christian Klämbt. 2020. “The Drosophila NCAM Homolog Fas2 Signals Independently of Adhesion.” Development 147 (2): dev181479.

Neville, Kathryn E, Tara M Finegan, Nicholas Lowe, Philip M Bellomio, Daxiang Na, and Dan T Bergstralh. 2023. “The Drosophila Mitotic Spindle Orientation Machinery Requires Activation, Not Just Localization.” EMBO Reports 24 (3): e56074. 10.15252/embr.202256074.

Olivieri, Daniel, Martina M Sykora, Ravi Sachidanandam, Karl Mechtler, and Julius Brennecke. 2010. “An in Vivo RNAi Assay Identifies Major Genetic and Cellular Requirements for Primary piRNA Biogenesis in Drosophila.” The EMBO Journal 29 (19): 3301–17.

Packard, Adam, Kylie Georgas, Odyssé Michos, Paul Riccio, Cristina Cebrian, Alexander N Combes, Adler Ju, et al. 2013. “Luminal Mitosis Drives Epithelial Cell Dispersal within the Branching Ureteric Bud.” Developmental Cell 27 (3): 319–30.

Resnik-Docampo, Martin, Christopher L. Koehler, Rebecca I. Clark, Joseph M. Schinaman, Vivien Sauer, Daniel M. Wong, Sophia Lewis, Cecilia D’Alterio, David W. Walker, and D. Leanne Jones. 2017. “Tricellular Junctions Regulate Intestinal Stem Cell Behaviour to Maintain Homeostasis.” Nature Cell Biology 19 (1): 52–59. 10.1038/ncb3454.

Robinson, James T, Helga Thorvaldsdottir, Douglass Turner, and Jill P Mesirov. 2023. “Igv.Js: An Embeddable JavaScript Implementation of the Integrative Genomics Viewer (IGV).” Bioinformatics 39 (1): btac830. 10.1093/bioinformatics/btac830.

Sauer, F C. 1937. “Some Factors in the Morphogenesis of Vertebrate Embryonic Epithelia.” Journal of Morphology, 1–19.

Spear, Philip C., and Carol A. Erickson. 2012. “Interkinetic Nuclear Migration: A Mysterious Process in Search of a Function.” Development, Growth & Differentiation 54 (3): 306–16. 10.1111/j.1440-169X.2012.01342.x.

Szafranski, Przemyslaw, and Scott Goode. 2007. “Basolateral Junctions Are Sufficient to Suppress Epithelial Invasion during Drosophila Oogenesis.” Developmental Dynamics : An Official Publication of the American Association of Anatomists 236 (2): 364–73.

Wei, Jun, Michael Hortsch, and Scott Goode. 2004. “Neuroglian Stabilizes Epithelial Structure during Drosophila Oogenesis.” Developmental Dynamics : An Official Publication of the American Association of Anatomists 230 (4): 800–808.

Williams, Scott E., and Kendall J. Lough. 2020. “Cell Biology: Pardon the Intrusion.” Current Biology 30(24): R1481–84. 10.1016/j.cub.2020.10.036.

Wilson, Tyler J, and Dan T Bergstralh. 2017. “Cell Reintegration: Stray Epithelial Cells Make Their Way Home.” BioEssays : News and Reviews in Molecular, Cellular and Developmental Biology 39(6): 1600248.

