## Supplementary Tables for "*Fas2^EB112^*: A Tale of Two Chromosomes"

**Table 1 – Reagents**

| **Reagent** | **Citation** | **Source** |
|---|---|---|
| rabbit anti-Centrosomin | Lucas & Raff, 2007 | Gift from Raff lab |
| rabbit anti-Inscuteable | Kraut et al, 1996 | Gift from Knoblich lab |
| rabbit anti-Nrg | This study | |
| mouse anti-Nrg (1B7) | Bieber 1989 | Hortsch lab |
| mouse anti-Dlg (4F3) | | DSHB 2/18/16 |
| mouse anti-Fas2 (1D4) | | DSHB 6/9/16 |
| Conjugated secondary antibodies | | ThermoFisher Scientific |
| mouse FITC-conjugated anti-α-tubulin (DM1A) | | Sigma Lot#114M4817V |
| Fluorescein Phalloidin | | Invitrogen: Lot# 2138394 |
| Alexa fluor 633 Phalloidin | | Invitrogen: Lot# 2274768 |
| Vectashield with DAPI | | Vector Labs H-1200 : Lot# ZK03217 |
| Pierce ECL Plus Substrate | | Thermo Fisher: Lot# RH23622 |
| Pierce protease inhibitor | | Thermo Fisher: Lot# SB2334961 |
| Amido Black | | Sigma: Lot# SLBR9882V |
| Sigmacote | | Sigma-Aldrich SL2 |
| Formaldehyde (37%) | | Thermo Fisher BP531 |
| Acetic Acid | | Thermo Fisher A507 |
| Poly-L-Lysine Coated Slides | | Electron Microscopy Sciences 6341001 |

**Table 2 - *Drosophila* Alleles and Transgenes**

| Mutant Alllele or Transgene | Citation | Stock Number or Lab of Origin |
|---|---|---|
| *Fas2*$^{EB112}$ FRT101 | | BDSC 36284 |
| *Fas2*$^{EB112}$ FRT19A | | Riechmann lab via Juan Manuel Gomez |
| *Nrg*$^{14}$ | | BDSC 36279 |
| Traffic jam-GAL4 | (Olivieri et al, 2010) | Lehmann lab |
| UAS-Fas3-RNAi TRiP.HMC06528 | (Perkins et al, 2015) | BDSC 77396 |
| UAS-Fas2-RNAi TRiP.HMS01098 | (Perkins et al, 2015) | BDSC 34084 |
| UAS-Nrg-RNAi TRiP.HMS01638 | (Perkins et al, 2015) | BDSC 367496 |
| Ubi-mRFP.nls, hsFLP, FRT19A | | BDSC 31418 |
| GFP, FRT101; hsflp | | Bilder lab |
