## Supplementary figures and images for "*Fas2^EB112^*: A Tale of Two Chromosomes"

### Supplementary Figure 1

#### SUPPLEMENTAL FIGURE 1 - Finegan *et al.*

**A**

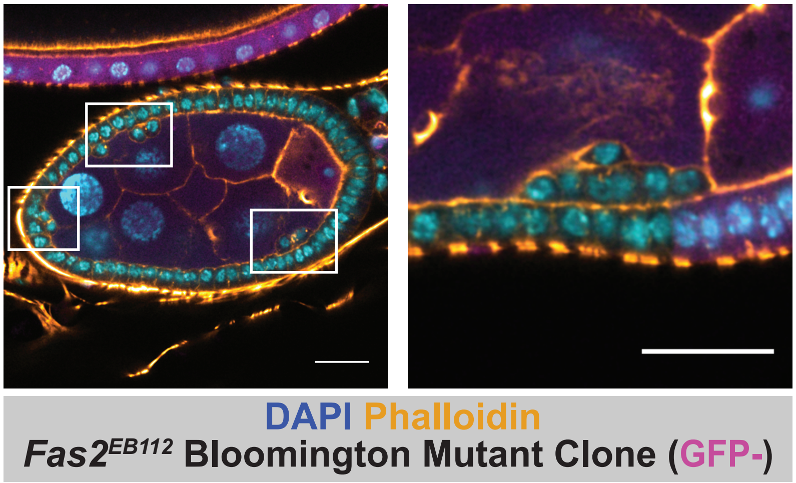


**B**

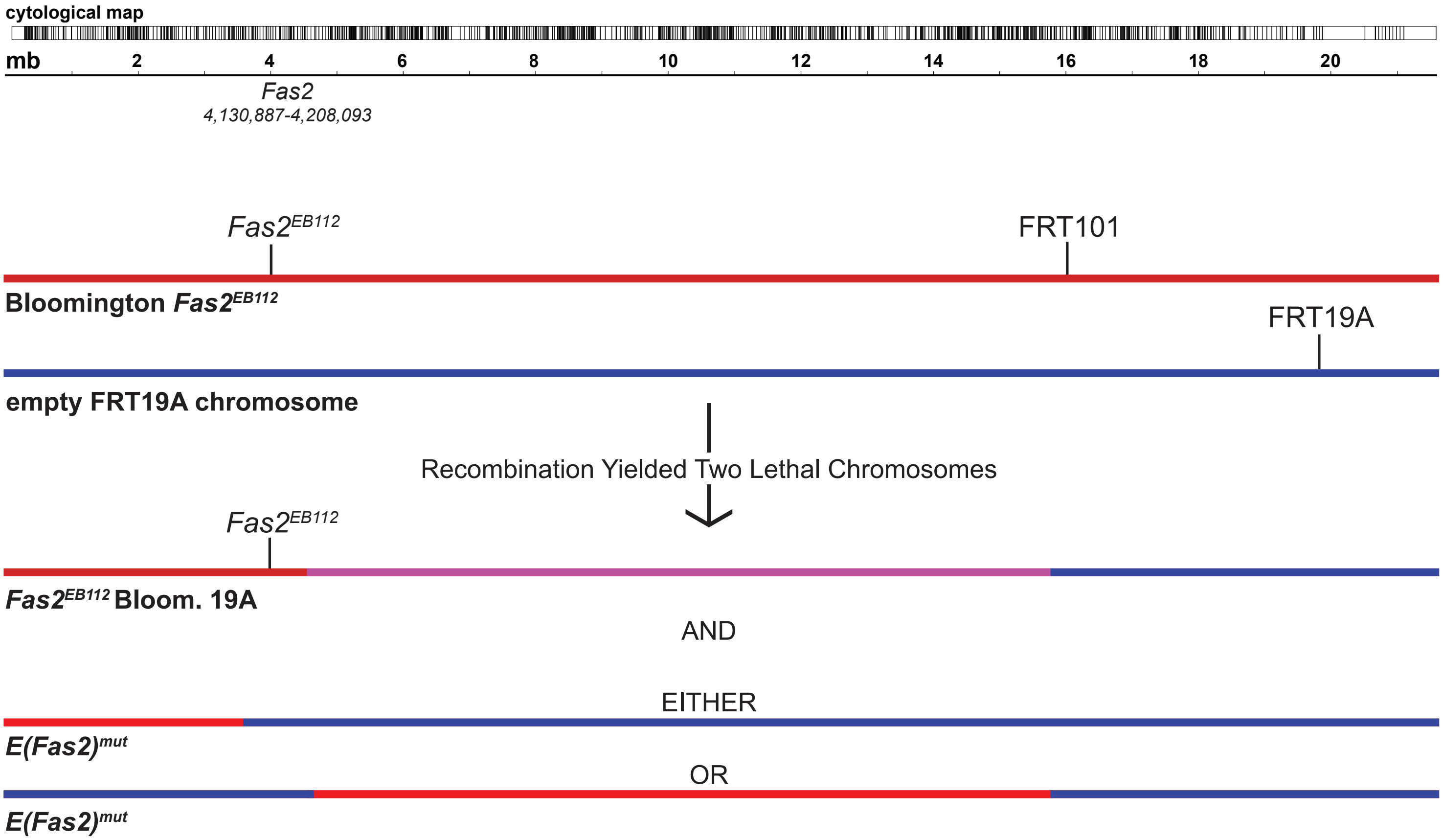

### Supplementary Figure 2

#### SUPPLEMENTAL FIGURE 2 - Finegan *et al.*

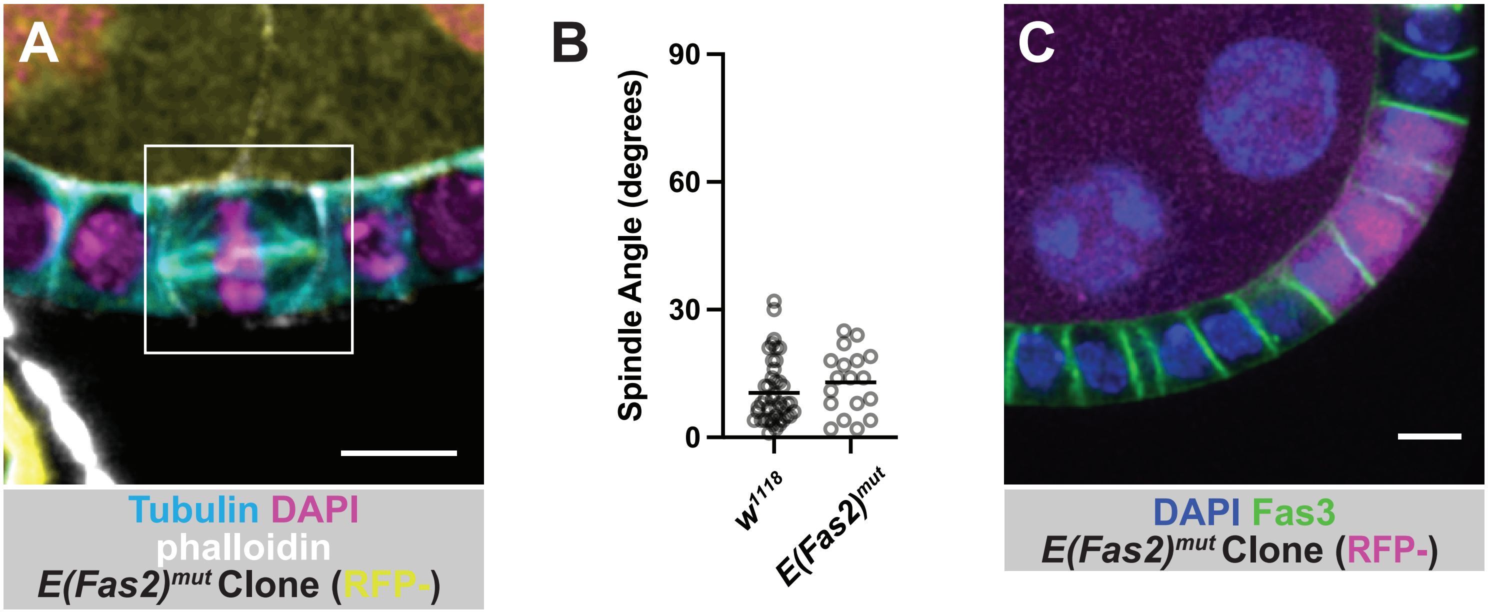

### Supplementary Figure 3

#### SUPPLEMENTAL FIGURE 3 Finegan *et al.*

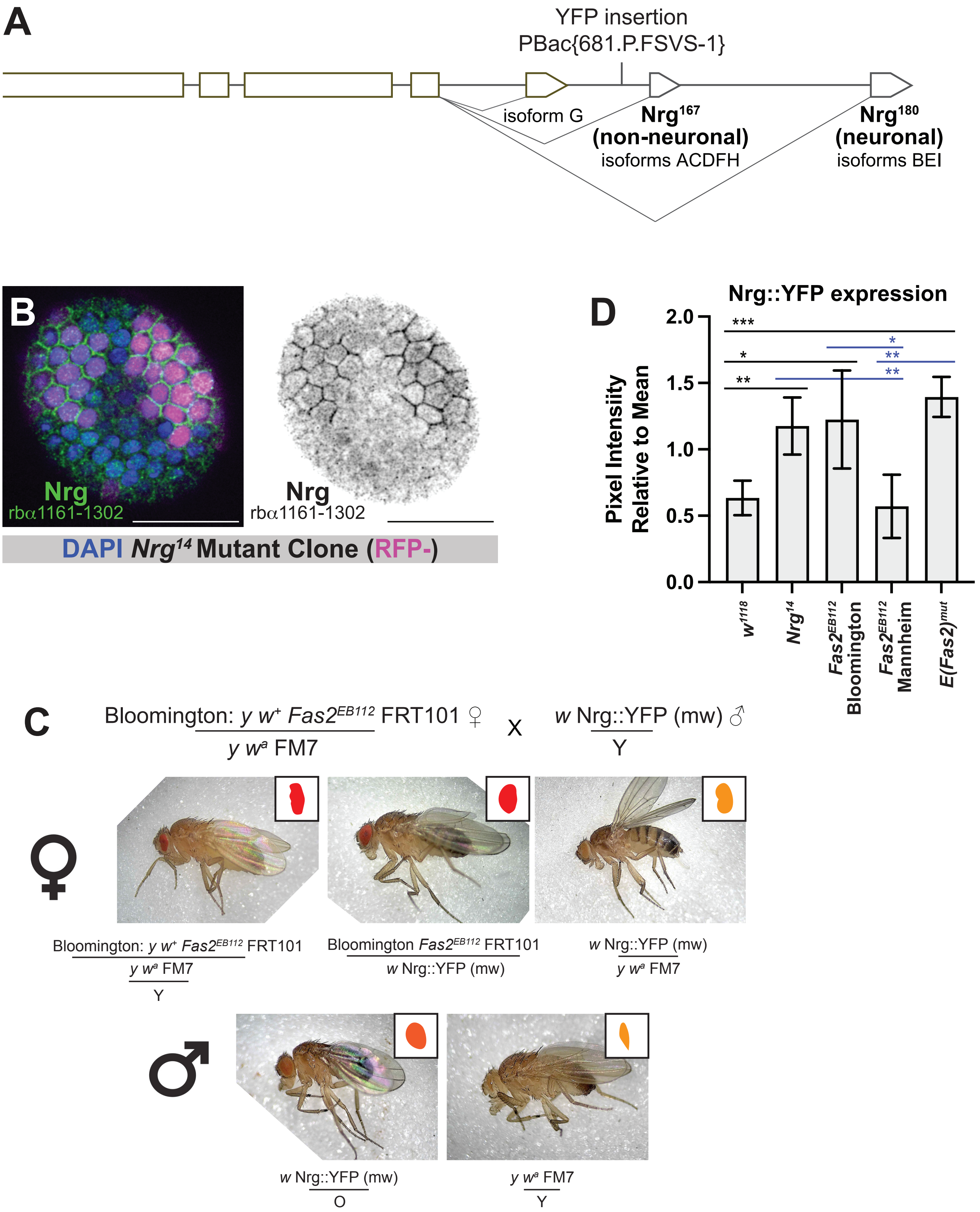

### Supplementary Figure 4

#### SUPPLEMENTAL FIGURE 4 - Finegan *et al.*

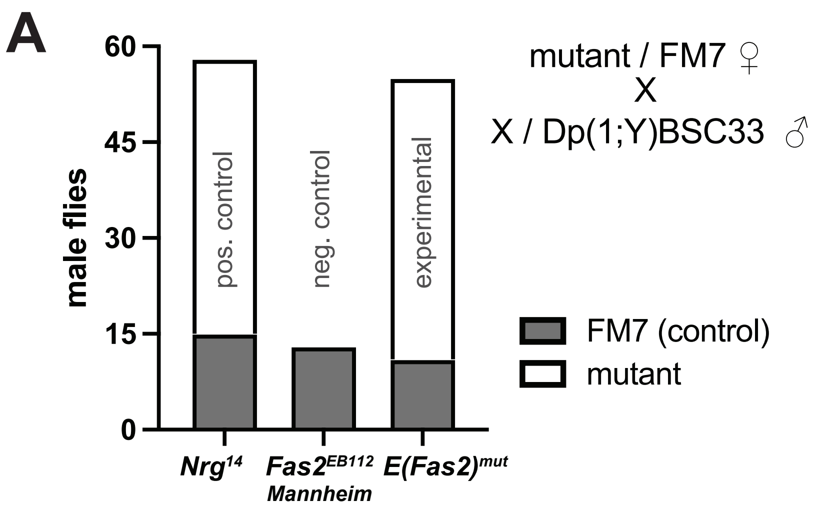


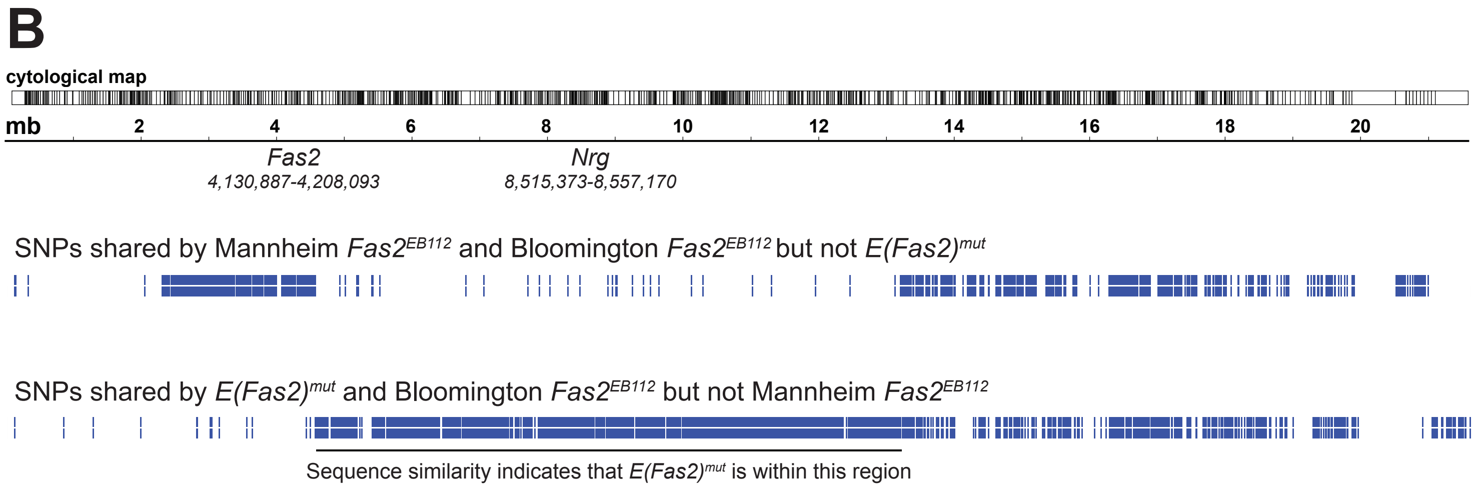
